# Metals alter membership but not diversity of a headwater stream microbiome

**DOI:** 10.1101/2020.04.30.071522

**Authors:** Brian A. Wolff, William H. Clements, Ed K. Hall

## Abstract

Metal contamination from mining or natural weathering is a common feature of surface waters in the American west. Traditionally, stream macroinvertebrate community metrics have been used for stream quality assessments. Advances in microbial analyses have created the potential for routine sampling of aquatic microbiomes as a tool to assess the quality of stream habitat. We sought to determine if microbiome diversity and membership were affected by metal contamination in a manner similar to what has been observed for stream macroinvertebrates, and if so, identify candidate microbial taxa to be used to indicate metal stress in stream ecosystems. We evaluated microbiome membership from sediments at multiple sites within the principal drainage of an EPA superfund site near the headwaters of the Upper Arkansas River, Leadville, CO. From each sample, we extracted DNA and sequenced the 16S rRNA gene amplicon on the Illumina MiSeq platform. We used the remaining sediments to simultaneously evaluate environmental metal concentrations. We also conducted an artificial stream mesocosm experiment using sediments collected from two of the observational study sites. The mesocosm experiment had a 2×2 factorial design: 1) location (upstream or downstream of contaminating tributary), and 2) treatment (metal exposure or control). We found no difference in diversity between upstream and downstream sites in the field. Similarly, diversity changed very little following experimental metal exposure. However, microbiome membership differed between upstream and downstream locations and experimental metal exposure changed microbiome membership in a manner that depended on origin of the sediments used in each mesocosm.

**Importance:** Our results suggest that microbiomes can be reliable indicators of ecosystem metal stress even when surface water chemistry and other metrics used to assess ecosystem health do not indicate ecosystem stress. Several results presented in this study are consistent with the idea that a microbial response to metals at the base of the food web may be affecting consumers one trophic level above. If effects of metals are mediated through shifts in the microbiome, then microbial metrics, as presented here, may aid in the assessment of stream ecosystems health.

## Introduction

Streams in the western United States are frequently impaired from elevated metal concentrations due to a combination of historical mining activities and to a lesser extent natural weathering processes. In Colorado, there are approximately 23,000 abandoned mines (1) resulting in approximately 23% of Colorado streams qualifying as impaired (2). One metric routinely used to evaluate stream water quality and ecosystem health is stream macroinvertebrate community composition. Various protocols that use macroinvertebrates continue to be standard practice for stream biomonitoring (3-5). Macroinvertebrates have a series of characteristics that have proven useful for stream bioassessments including ubiquity, high diversity, restricted ranges, short generation times, small size, and are important food sources for aquatic and terrestrial consumers alike (6). Because microorganisms have many of these same characteristics and because analyses of microbiome characteristics have become more routine, we investigated whether microbiomes had dynamic responses to metal exposure in metal-contaminated ecosystems that would make them an appropriate indicator of stream ecosystem health.

We hypothesized that microbiomes may potentially be better indicators of water quality than macroinvertebrates because they are even more ubiquitous and dynamic and thus may report even subtler differences in water quality. For instance, typical bacterial generation times (i.e., doubling time) occur over hours or days (7) compared to weeks to months for macroinvertebrates (8). The spatial scale at which microbiomes operate is also much smaller than for stream macroinvertebrates, creating potential to identify small pockets of contamination in heterogeneous stream ecosystems. We now know that microbial biofilms are formed by complex, non-random assemblages of algae, bacteria, and fungi (9) and that these diverse microbiomes can be shaped by physical properties like stream velocity (10) and substrate type (11) as well as chemical properties such as pH (12, 13). It is also clear that metals affect the function of microbiomes, including evidence for metals decreasing stream nitrification (14), and reducing rates of microbial respiration (15). Metals also affect microbiome membership, including evidence where specific sub-phyla increased (γ-proteobacteria) or decreased (β- proteobacteria) with metal exposure (16).

To test the potential for microbiomes to act as indicators of metal contamination we evaluated the stream microbiome of the Upper Arkansas River near Leadville, Colorado, USA. The Upper Arkansas River has been impaired by metal pollution due to historical mining since the mid-1800s (17). By the late 1990s, implementation of water treatment facilities and removal of floodplain mine tailings resulted in significant improvements in water quality including decreased dissolved metals – principally cadmium (Cd), copper (Cu), and zinc (Zn) – downstream from where California Gulch enters the mainstem of the Upper Arkansas River (17). Despite improved water quality, macroinvertebrate community membership has remained different between upstream reference sites and sites downstream of California Gulch (17). Although species richness has remained similar between upstream and downstream locations, community membership has continued to differ among sites (18).

To assess how microbiomes were affected by metal exposure in the Upper Arkansas River we chose to focus on the bacterial component of the stream microbiome because: A) sediment biofilms are primarily composed of bacterial biomass (from 90-99%) (19, 20), B) many stream macroinvertebrates spend significant portion of their lifecycle grazing on biofilm in sediments (21) so changes in microbiome may have effects on higher trophic levels, and C) 16S rRNA gene sequences have better developed sequence libraries compared to analogous phylogenetic markers for other groups, such as the 18S rRNA gene for eukaryotic microbes (22). We collected samples at locations upstream and downstream of California Gulch during both Spring and Fall seasons. From each sample, we used 16S rRNA amplicon sequencing of the sediment biofilms on the Arkansas River to determine how metals influenced microbiome diversity and membership in the Upper Arkansas River. We complemented our field observations with experiments that exposed microbial communities sampled from both upstream and downstream of California Gulch to elevated metal concentrations. The purpose of this study was to examine: (1) how microbiome diversity and membership differed in an ecosystem that has elevated metal levels, (2) if these differences could be attributed to exposure to metals and (3) if certain microbiome genera were consistently enriched or depleted in response to metal exposure and therefore may be candidates for indicators of stream water quality. The last goal is an important step toward identifying mechanistic responses of individual bacteria and aid in the development in using certain groups as sensitive “indicators” of metals stress.

## Methods

### Study Site

We conducted our observational study on the Upper Arkansas River, located near the town of Leadville, approximately 100 km west of Denver, Colorado. This area of the Upper Arkansas has been monitored since 1989 and the site conditions are well characterized in previous studies (17). Briefly, this area is approximately 2,820 meters above sea level, and typically receives ∼30 cm of precipitation annually. The Arkansas River has a snowmelt driven hydrograph, with peak discharges in May or June, normally reaching base flow conditions by September. Variable run-off alters streamflow and contributions of solutes (including metals) from the watershed, resulting in higher metal concentrations recorded during Spring (i.e., during snowmelt runoff) compared to the Fall (i.e., at baseflow) (17). In the reach of the Upper Arkansas we evaluated the stream substrate was primarily composed of medium to large cobble in a matrix of gravel and sand. Most riparian vegetation was composed of sagebrush (*Artemisia* spp.), grasses, and willow (*Salix* spp.) trees.

### Observational Study

We sampled sediment bacteria communities in the main stem of the Arkansas River at three locations: 2 sites upstream (AR1 and AR2), 4 sites downstream (AR3, AR4, AR4G, and AR5), and 1 site within the principal metal contributing tributary, California Gulch (Figure 1). At each site we collected samples to be analyzed for metal concentration and 16S amplicon sequencing in Spring (first week of May) and Fall (first week of October) of 2017. For each sampling event we collected four independent sediment samples in riffle habitat (with a water column depth of ca. 0.25 m) at each of the 7 sites. For each sample, we removed a large cobble (∼ 0.3 m diameter) and scooped underlying sediments into separate 50 ml Falcon™ tubes.

**Figure 1.**
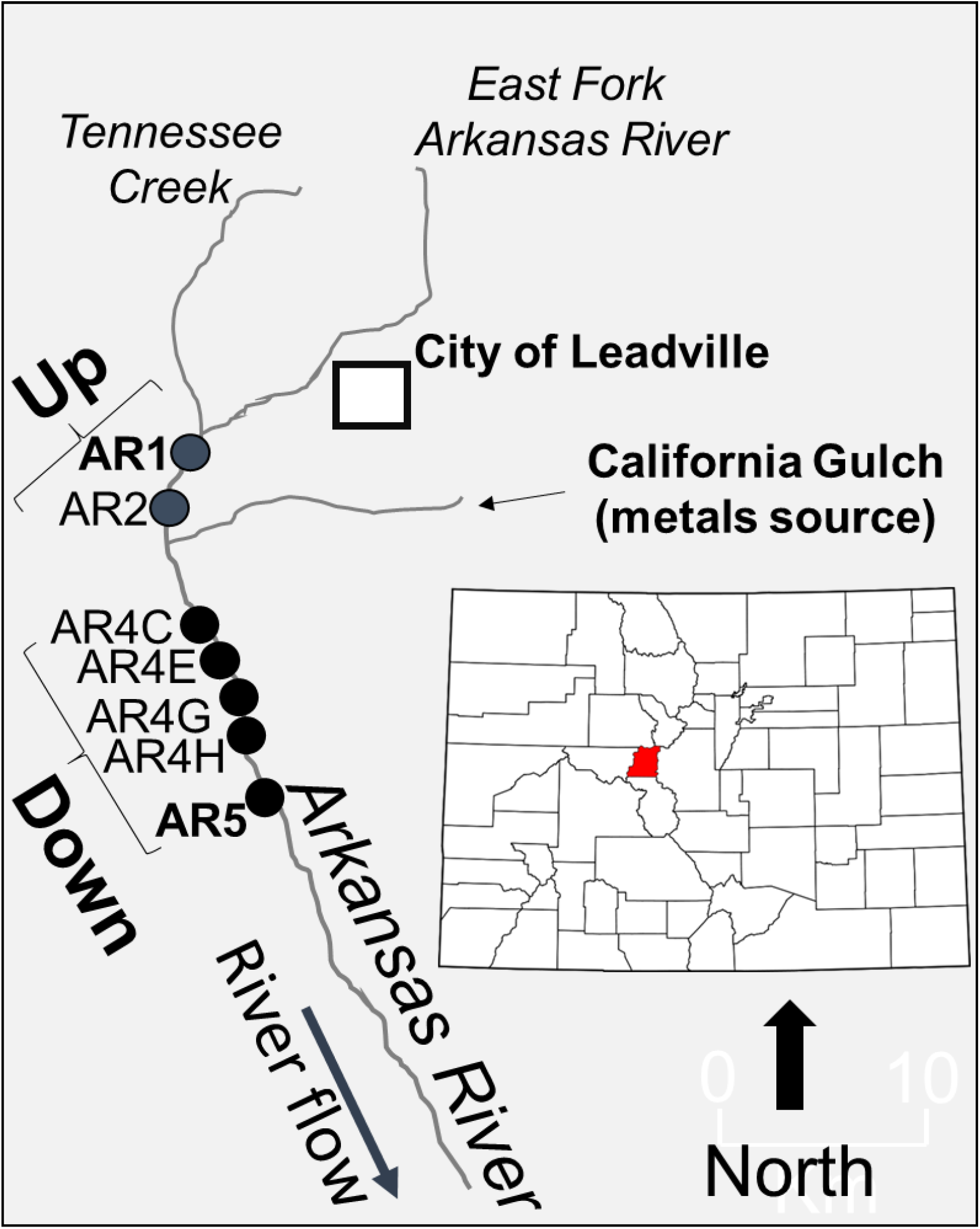
Map of the study area in the Upper Arkansas River, Colorado, USA.

### Experimental Mesocosm

To more explicitly test the effects of metals on stream microbiomes, we designed an artificial mesocosm experiment using samples from an upstream and downstream location. Specifically, we tested if experimentally manipulated metal concentrations would result in similar effects on the microbiome as seen from the metal gradient in the Upper Arkansas River. The observational study was conducted in Spring and Fall, however the mesocosm experiments were conducted only in the Fall because we were primarily interested in the differences in communities under stable conditions (e.g., base flow) and less by short-term seasonal effects from spring snowmelt. The design and parameters of the mesocosm experiments have been described elsewhere (23). Briefly, biofilms for the experiments were collected by placing plastic trays containing clean (scrubbed and air-dried) cobble in the river for 31 days (09/05/2017 – 10/06/2017) allowing microbial biofilms to colonize the cobble in each tray. Trays were deployed at one reference site upstream of California Gulch (AR1; hereafter “Upstream”) and one site downstream of California Gulch (AR5, hereafter “Downstream”). Upon retrieval, 4 colonized trays were collected from each site and placed into individual coolers filled with ambient stream water then immediately transported to CSU’s Stream Research Laboratory (∼3 hours from the sampling site). The 4 trays from each cooler were then placed into an individual experimental “racetrack” stream which after an equilibration period (∼ 24 hours) were randomly assigned to 2 treatments: metals or no metals (control).

Each artificial stream received source water from the hypolimnion of a mesotrophic reservoir (Horsetooth Reservoir) that was delivered at a rate of 1.0 L min^-1^, resulting in a residence time of approximately 20 minutes for each mesocosm. Characteristics of the source water (e.g., pH, conductivity, temperature, dissolved oxygen) were typical of non-polluted mountain streams in Colorado (24). We implemented a 2 × 2 (location X treatment) factorial experimental design: 1) control-upstream, 2) control-downstream, 3) metals-upstream, and 4) metals-downstream. Each control and treatment were replicated four times for a total of 16 experimental streams. We started metal additions after a 24 hr acclimation period. Peristaltic pumps delivered stock solutions of a metal mixture from a 20 L concentrated carboy at a rate of 10 ml min^-1^ to obtain a targeted concentration of 25 µg L^-1^ Cu and 650 µg L^-1^ Zn for each treatment. Paddlewheels provided a constant flow of 0.35 m s^-1^ to each mesocosm. Metals concentrated in the carboys were refreshed daily during the 10-day experiment. We checked water and peristaltic pump flows twice daily to ensure consistent delivery of metal solutions among treatments. We measured ambient metals concentrations from each mesocosm by filtering (0.45 µm) 15 mL water samples on Day 2, Day 4, and Day 10 of the experiment. On Day 10, all trays from each stream were collected, sieved (350 µm) into a clean, plastic bucket. Buckets were then decanted and the remaining material (e.g., sediments and periphyton floc) were transferred into 50 ml Falcon tubes and frozen at −80 °C until DNA extraction and metals analysis.

### DNA preparation and 16S rRNA amplicon sequencing

We extracted DNA from each sample with a MoBio PowerSoil® DNA Isolation Kit using standard protocols. The 16S rRNA gene (V4 region) was amplified using primers 515F and 806R universal primers with the forward primer barcoded following the Earth Microbiome Project protocols (25). The forward primer 515F included the unique sample barcode following Parada et al. (26), and both primers included degeneracies as described in Parada et al. (26) and Apprill et al. (27). For each sample, we ran a 50 µL PCR reaction using an Invitrogen PlatinumTM Hot Start PCR Master Mix with 10 µL of DNA. The PCR product was quantified and then pooled into a single pool in equimolar concentrations and cleaned using a MinElute® PCR Purification kit. Cleaned, pooled DNA was sequenced with a MiSeq reagent v2 500 cycle kit on the Illumina MiSeq platform at the Colorado State University Next Generation Sequencing Core facility.

Sequence reads were analyzed using MOTHUR (28) and OTUs counts defined at a 97% similarity of the sequence using the OptiClust algorithm. Generated OTUs were then aligned to a SILVA reference file (29). After sequences were processed through the MOTHUR pipeline, we then imported the data in R studio (30) for statistical analyses and visualization. Within R, subsequent analyses were performed utilizing the package Phyloseq. Sequences were pre-processed to remove Operational Taxonomic Units (OTUs) that were not counted at least 3 times in 20% of the samples.

For most analyses, raw OTU counts were transformed to relative abundances within each sample to reduce issues that can arise from count data from samples with varying library size. However, for DESeq2 analyses, we did not normalize count data to relative abundances because DESeq2 algorithms require raw sequence count data inputs. We also aggregated all OTUs that shared the same genera before performing DESeq2 analysis. We visualized DESeq2 results with Log2-fold change plots analyses. All sequences have been uploaded to the Sequence Read Archive (SRA) (31) and can be accessed from the NCBI BioProject accession number PRJNA628700.

### Metals preparation

We measured metal concentrations using material remaining from sediment samples that had previously been sub-sampled for DNA preparation. We dried sediments in a drying oven at 60 °C for at least 24 hours with periodic weighing of each sample until no more mass was lost and the sample remained at a constant weight. A small amount of sediment (0.14 – 0.25 g) was then weighed and transferred into 15 mL Falcon® tubes. Next, 1 mL trace-element grade nitric acid (HNO_3_) was added to each sample. Samples were vortexed and then placed in a hot water bath at 90 °C for 4 – 6 hours. Samples were then cooled outside of the hot water bath and ca. 0.2 mL hydrogen peroxide (H_2_O_2_) was then added. Samples were vortexed again and then placed back in the hot water bath for an additional 4 – 6 hours. After this period, samples were cooled, and 8.8 mL of Milli-Q water was added to ensure that all samples were diluted to a total of 10 mL. Samples were vortexed for a final time, centrifuged at 2500 rpms for 5 minutes, and the supernatant was extracted into clean 15 mL falcon tubes for quantification of metal concentration. We used dry weight and dilution volume (10 mL) to calculate the concentration of metals in each sample (µg g^-1^). Metal concentrations were quantified using a flame Atomic Absorption Spectrometer at the Colorado State University’s Aquatic Ecotoxicology Laboratory. From each sample we measured copper (Cu), cadmium (Cd; observational samples only), and zinc (Zn). These metals have been determined from previous studies to be the principal metals contaminating the Upper Arkansas River (17).

### Biomonitoring statistics

We estimated microbiome alpha diversity using: 1) richness from the number of unique OTUs present in each sample; and 2) the Shannon Index (32) that accounts for both richness and evenness from the distribution of those OTUs. To assess the effect of metals on membership we grouped upstream sites (AR1 and AR2) together (subsequently referred to as “Upstream”), and we grouped the four downstream sites (AR3, AR4, AR4G, and AR5) together (subsequently referred to as “Downstream”). To test for differences in microbiome membership between upstream and downstream communities we used a Bray-Curtis Similarity Index and visualized the similarities in community membership using Principle Coordinates Analysis (PCoA plot). To determine if clusters from each location/season were statistically different from each other we used a PERMANOVA model. For the *in-situ* Arkansas River bacterial communities, we tested the effect of location (upstream vs. downstream), season (Spring vs. Fall), and their interaction. PERMANOVA tests were also used for pairwise comparisons (e.g., Upstream-Spring vs. Downstream-Spring; Upstream-Fall vs. Downstream-Fall, etc.).

After evaluating whole community membership differences, we used log2-plots to visualize what, if any, microbiome genera were significantly enriched or depleted in the upstream or downstream locations. We used a very low alpha value (α ≤ 0.0005) to protect against type-I error when identifying genera that are differentially enriched between upstream and downstream locations. If certain genera were more enriched at downstream locations (positive log2-fold value) we considered them as potential indicators of metal-tolerant bacteria. Similarly, genera more enriched at upstream sites (negative log2-fold value) we considered to be indicators of metal-sensitive bacteria. To aid in comparison with mesocosm results, we only included samples sourced directly from AR1 (upstream) and AR5 (downstream) locations for Log2-fold plots. *Mesocosm Statistics* For the mesocosm experiment samples, we used a 2×2 factorial design that tested the effect of metals treatment (control *vs*. treatment), and location (upstream *vs*. downstream). We used many of the same statistical and visual analyses as described previously for the observational study. One notable difference is that for mesocosm log2-fold plots, we compared metal-treated groups (positive log2-fold values) to control groups (negative log2-fold values) separately by each location.

## Results

### Observational Study

To assess the effect of metals on the Upper Arkansas River microbiome during the Spring and Fall of 2017 we analyzed *in situ* pH, sediment metal concentration, and 16S rRNA amplicons from California Gulch and sites upstream and downstream of California Gulch in the main stem of the Upper Arkansas River. For all sampling locations during both seasons California Gulch had the highest sediment metal concentrations for all three metals, followed by downstream sites, with upstream sites consistently having the lowest sediment metal concentrations (Table 1). At downstream sites there was a statistically significant increase of Cu and Zn in the Spring compared to Fall, but not for Cd (Table 1). However, we did not observe statistically different metal concentrations between Fall and Spring for any of the metals at the upstream sites (consistently low) or in California Gulch (consistently high, Table 1).

**Table 1.** Descriptive statistics for the observational study in the Upper Arkansas River. Different letters refer to a statistically significant difference (α = 0.05) among locations (upstream, California Gulch [CalGulch], and downstream samples) in the Spring (lower case) and in the Fall (Upper case). Asterisks refer to statistically significant difference (α = 0.05) between Spring and Fall for each location.

| Location | Season | n | pH |  | Sediment Cd |  | Sediment Cu |  | Sediment Zn |  | Observed OTUs |  | Shannon Index |  |
| --- | --- | --- | --- | --- | --- | --- | --- | --- | --- | --- | --- | --- | --- | --- |
|  |  |  | Mean | StdDev | Mean | StdDev | Mean | StdDev | Mean | StdDev | Mean | StdDev | Mean | StdDev |
| Upstream | Spring | 8 | 7.63 a* | 0.21 | 1.59 a | 0.41 | 4.29 a | 1.15 | 220.24 a | 92.16 | 4825.38 a | 1131.69 | 6.56 a | 0.35 |
| CalGulch | Spring | 4 | 7.12 b* | - | 23.66 b | 10.15 | 331.30 b | 113.47 | 6300.8 b | 1673.60 | 2414.00 b | 1402.84 | 5.84 b | 0.25 |
| Downstream | Spring | 13 | 7.41 c* | 0.05 | 6.29 c | 4.09 | 53.56 c* | 69.58 | 1145.10 c* | 590.00 | 4095.31 a* | 1148.54 | 6.49 a | 0.33 |
| Upstream | Fall | 8 | 7.75 A** | 0.53 | 1.74 A | 0.58 | 4.19 A | 2.28 | 167.17 A | 111.22 | 5288.50 A | 1010.73 | 6.70 A | 0.23 |
| CalGulch | Fall | 4 | 7.50 B** | - | 33.45 B | 16.58 | 382.49 B | 174.56 | 6488.64 B | 1059.45 | 2729.50 B | 388.95 | 5.65 B | 0.06 |
| Downstream | Fall | 14 | 7.49 B** | 0.07 | 5.14 C | 2.81 | 18.13 C** | 8.24 | 767.12 C** | 395.35 | 5033.29 A** | 826.92 | 6.49 A | 0.41 |

We also examined surface water pH at the time of sediment sample collection since the addition of metals can lower pH of the receiving waters, and more importantly, lower pH can make metals more bioavailable to aquatic organisms. While differences in pH among locations were minimal (they ranged between 7.12 – 7.75), in general, samples with higher metal concentrations had lower pH (Table 1). There was also a statistically significant season by location interaction for pH (p=0.002). Spring pH values were different among all locations with upstream sites having the highest pH values followed by downstream sites, and then California Gulch. In the Fall, pH at upstream sites was significantly higher than California Gulch and all downstream sites, but there was no statistical difference between California Gulch and downstream locations (Table 1). For all locations, pH was lower in the Spring than in the Fall.

At each site we assessed the 16S rRNA amplicons from all sediment samples. Richness (i.e., number of observed OTUs) was significantly different among locations and between seasons (Table 1). California Gulch had the lowest richness among all sites in both Spring and Fall, however upstream and downstream locations had similar richness in both seasons (Table 1). At downstream locations, richness was significantly lower (p=0.0365) in the Spring than the Fall, but we observed no significant difference in microbiome richness between seasons at the upstream sites or in California Gulch (Table 1). Shannon Index values were lowest at California Gulch in both Spring and Fall (Table 1), and there was no statistical difference diversity between upstream and downstream locations (Table 1). However, unlike richness results, there was no difference in Shannon Index values between seasons, or any significant season by location interactions (Table 1).

We also evaluated microbiome membership to determine if metals altered the composition of the microbiome (i.e., membership) even though indices of alpha diversity may not have differed. We found that microbiome membership was significantly different among locations (p=0.001) and between seasons (p=0.015). The membership of the California Gulch microbiome was different than the membership of the microbiome at upstream and downstream locations for all sampling dates (Figure 2). Because the difference in membership between California Gulch and either location in the mainstem of the Arkansas River was so pronounced we also performed a PERMANOVA that excluded California Gulch from the dataset to focus on differences between upstream and downstream sites. We found a significant difference in membership (p=0.003) between locations and a significant within site difference between seasons (p=0.016). However, the season by location (i.e., upstream vs. downstream) effect was not significant (p=0.670), suggesting that microbiome membership differed between seasons, but changes occurred at both locations. A constrained analysis of principles coordinates (CAP) illustrated that differences in microbiome membership between locations were primarily driven by higher Cu and Zn sediment concentrations in downstream sites, which also likely reduced pH at the downstream sites (Figure 3).

**Figure 2.**
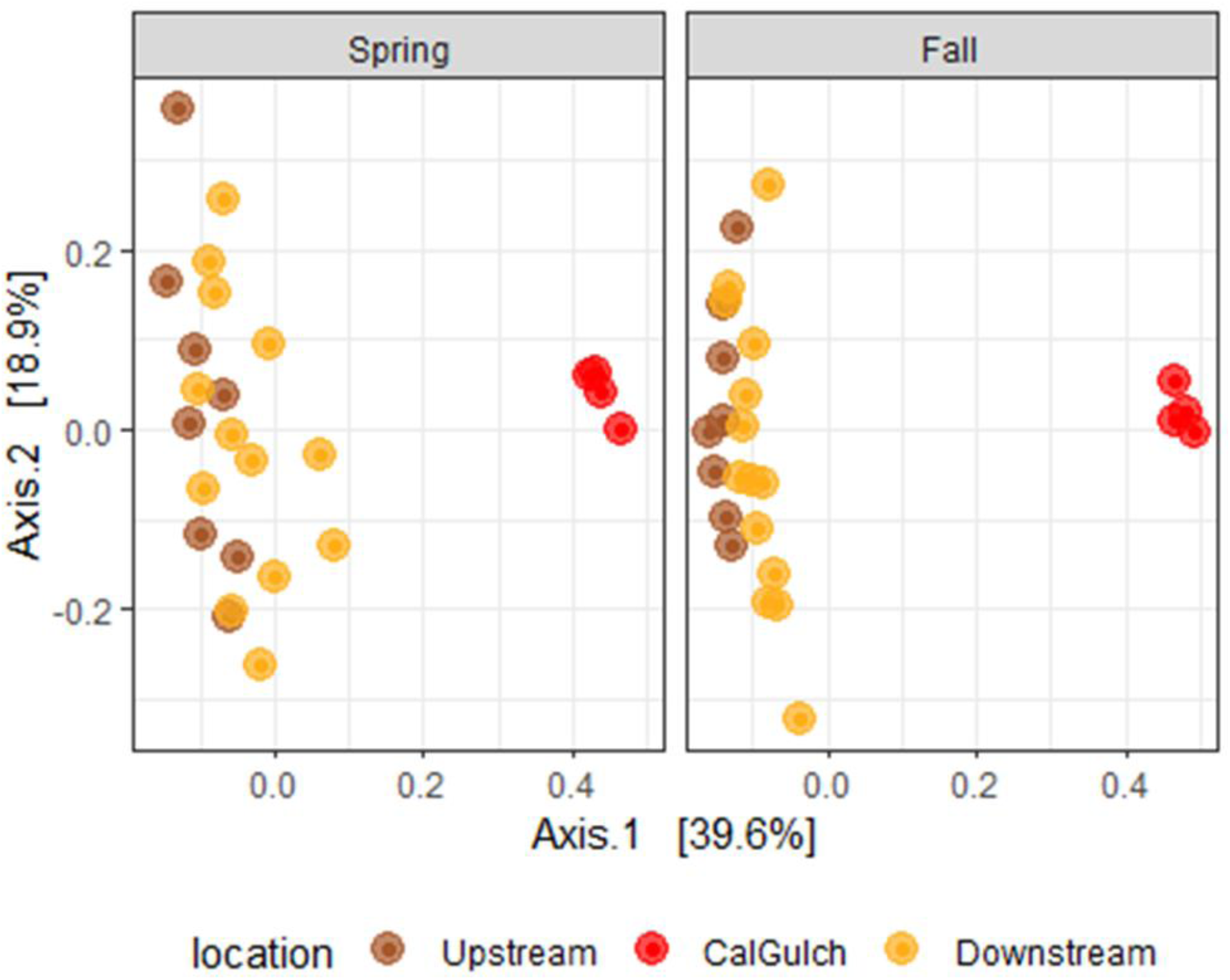
Principle Coordinates Analysis (PCoA) of microbiomes among locations both with California Gulch (CalGulch).

**Figure 3.**
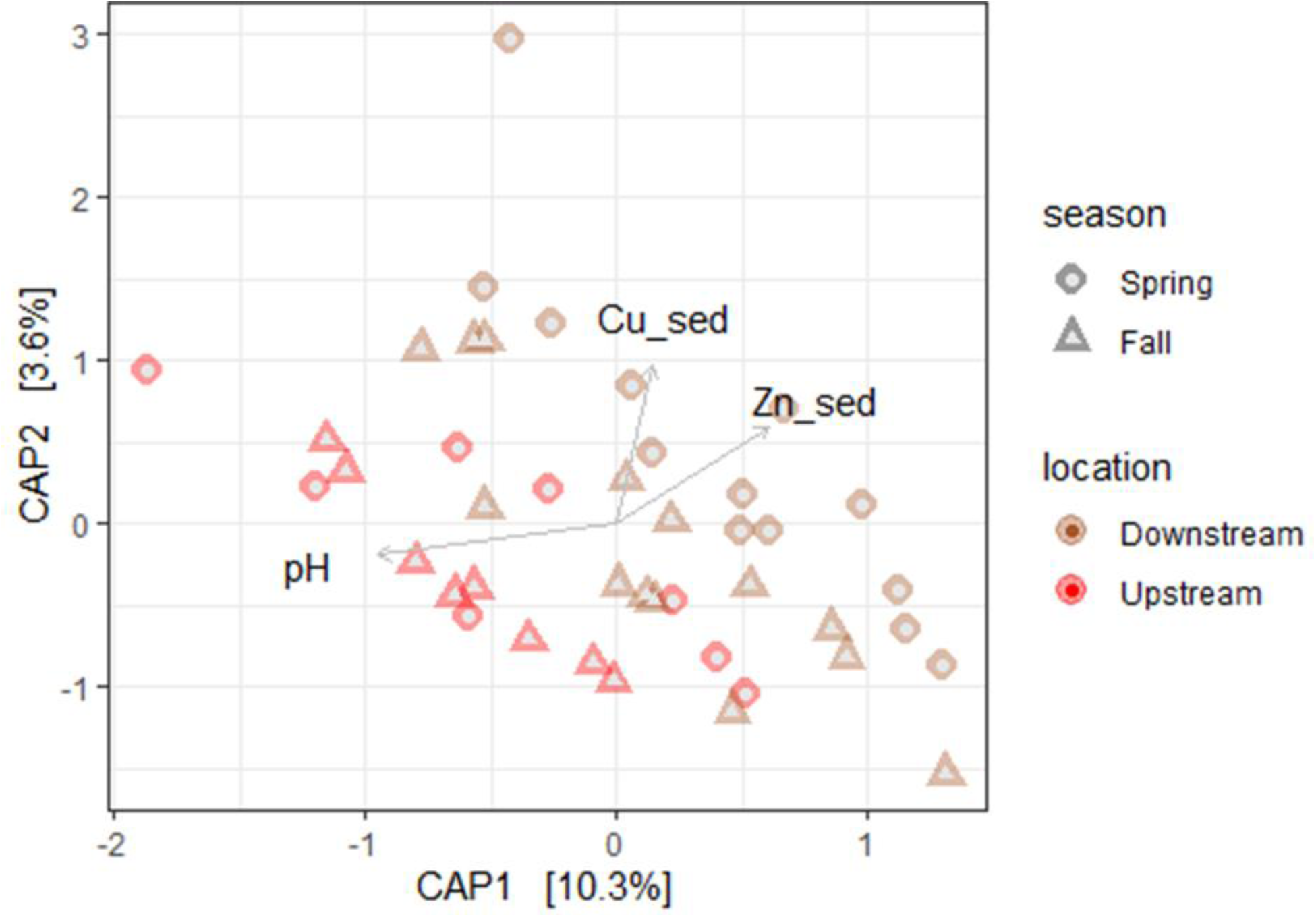
Constrained Analysis of Principles coordinates (CAP) analysis without California Gulch included.

### Mesocosm Study

We did not find significant effects of location, metals, or an interaction between location and metals, on richness in any of the mesocosms after the 10-day exposure period (Table 2). However, Shannon Index values were consistently higher for the microbiomes sourced from upstream sites compared to the microbiomes sourced from downstream sites for both the control and metal enriched treatments. When we assessed the effect of the metal treatment on diversity within a location, we found that Shannon Index values were significantly lower in the metal-treated samples for the downstream location, but we did not see a similar change in diversity in response to metals for the upstream location (Table 2).

**Table 2.** Descriptive statistics for the mesocosm study. Different letters refer to a statistically significant difference (α = 0.05) between upstream and downstream control samples (lower case), and between upstream and downstream metal-treated samples (Upper case). Asterisks refer to statistically significant difference (α = 0.05) between control and metal-treated samples for each location.

| Location | Season | Treatment | n | Water Cu (µg/L) |  | Water Zn (µg/L) |  | Sediment Cu (µg/g dry) |  | Sediment Zn (µg/g dry) |  | Observed OTUs |  | Shannon Index |  |
| --- | --- | --- | --- | --- | --- | --- | --- | --- | --- | --- | --- | --- | --- | --- | --- |
|  |  |  |  | Mean | StdDev | Mean | StdDev | Mean | StdDev | Mean | StdDev | Mean | StdDev | Mean | StdDev |
| Upstream | Fall | Control | 4 | 2.35 a | 2.65 | 0.05 a | 0.17 | 43.07 a | 17.85 | 1311.91 a | 229.62 | 4715.50 a | 2706.24 | 6.78 a | 0.12 |
| Downstream | Fall | Control | 4 | 1.33 a | 0.92 | 0.23 a | 0.77 | 75.93 b | 19.63 | 1600.16 b | 36.47 | 3728.50 a | 599.13 | 6.30 b* | 0.01 |
| Upstream | Fall | Metals | 4 | 17.21 A | 6.31 | 588.85 A | 57.08 | 162.71 A | 68.09 | 2451.97 A | 645.75 | 5668.25 A | 3493.11 | 6.72 A | 0.11 |
| Downstream | Fall | Metals | 4 | 18.04 A | 4.94 | 613.53 A | 49.32 | 656.28 B | 161.64 | 7280.31 B | 1578.52 | 3608.00 A | 1007.13 | 6.11 A** | 0.13 |

Interestingly, in the metal-treated samples more Cu and Zn were retained in the downstream sediments compared to the upstream sediments over the course of the experiment (Table 2). Although, downstream sediments likely started with greater metal concentrations (inferred from the observational study), we did not see a similar difference in concentrations in the control samples between upstream and downstream sediments suggesting that the metal-treated sediments retained metals during the course of the experiment. This also mirrored the response of the stream microbiome, where the effect of metals depended on the location from which sediments were sourced. Over the course of the incubation, microbiomes from the downstream site showed a more pronounced change in membership to metal exposure than the upstream microbiome (Figure 4). This result was supported by a PERMANOVA that identified significant differences in membership between location (p=0.001), treatment (p=0.001), and the location by treatment interaction term (p=0.007).

**Figure 4.**
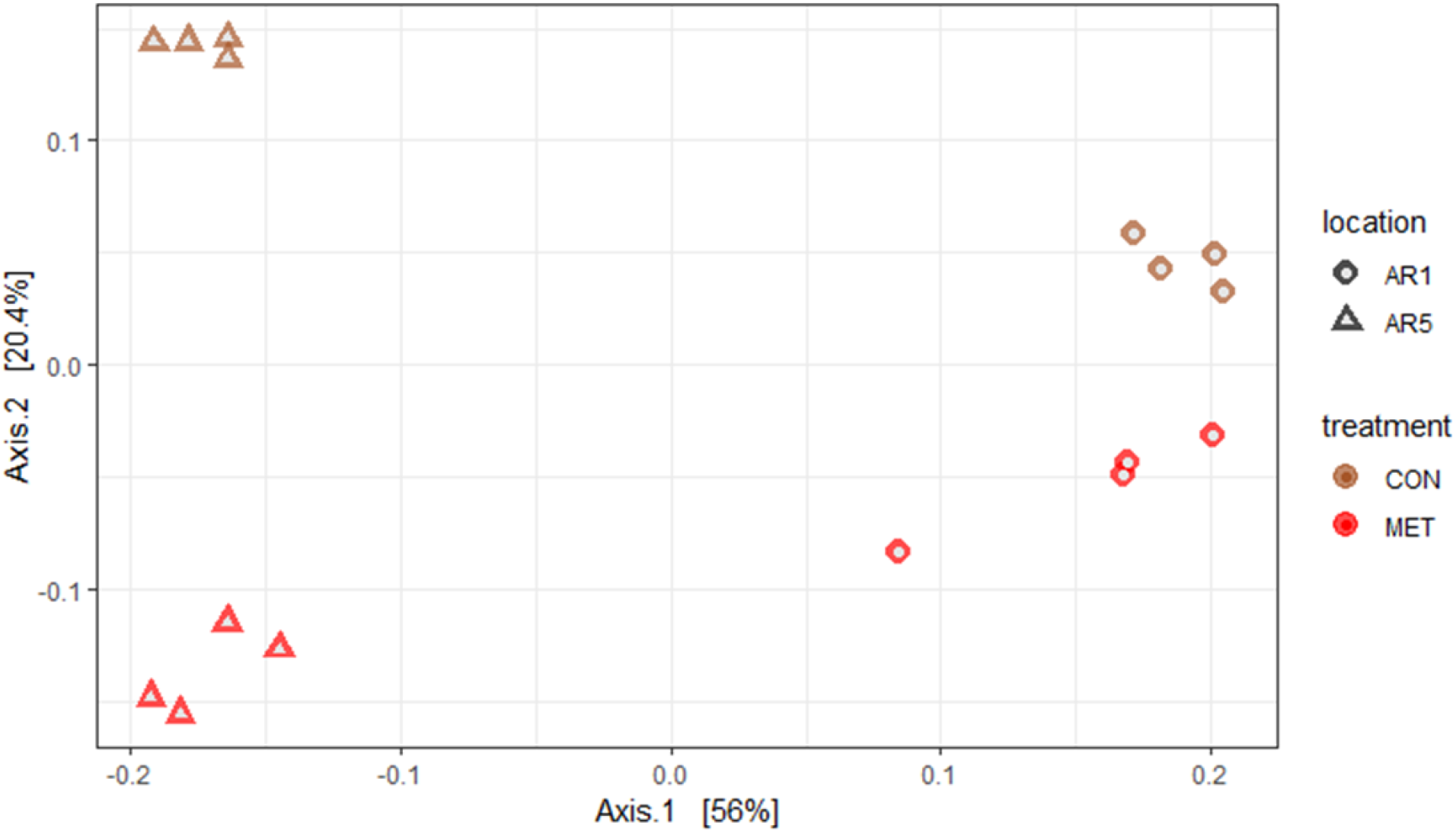
Principle Coordinates Analysis (PCoA) of community membership of upstream (circles) vs. downstream (triangles) sediment samples. Samples treated with metals are in red and non-treated controls are in brown.

### Genera-level responses between observational and mesocosm studies

In order to assess if microbiome membership was altered by the presence of metals similarly in our observational and experimental studies, we used Log2-fold plots to evaluate changes in genera (all OTUs identified to a common genus were aggregated) among the two components of the study. From the observational study we focused on the upstream (AR1) and downstream (AR5) sites that were used to seed the mesocosm experiments. The downstream site was significantly (p≤0.0005) enriched in genera from Cyanobacteria and Verrucomicrobia relative to upstream site (Figure 5). The upstream site had enriched genera from Latescibacteria, Acidobacteria, Proteobacteria, and Rokubacteria relative to the downstream site (Figure 5).

**Figure 5.**
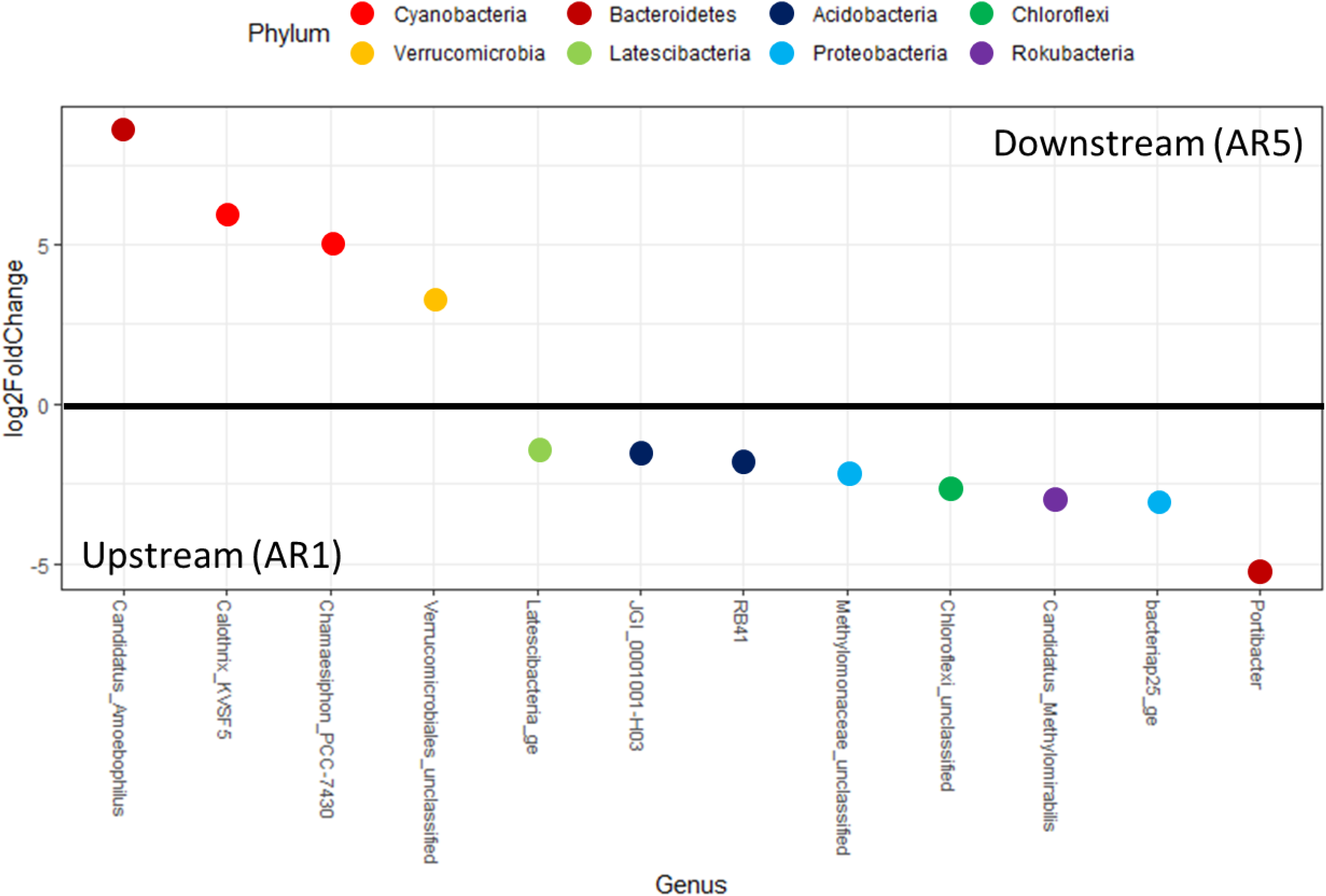
Log_2_-fold change plots of genera between upstream (AR1) and downstream (AR5) samples. Samples with a positive value are more enriched at upstream sites, and negative values more enriched at downstream sites. The color of each dot is the phylogenetic Order.

Microbiomes from each location had genera that were enriched in the metal-treated mesocosms compared to the control treatments. Both locations were enriched in genera from Bacteroidetes in the metal treatment compared to control mesocosms (Figure 6). Conversely, genera from Patescibacteria, Planctomycetes, Acidobacteria, Armatimonadetes, and Cyanobacteria were significantly enriched in control mesocosms (Figure 6). Some genera from Proteobacteria were significantly enriched metal-treated streams, while other genera from the same phylum were significantly enriched in control mesocosms.

**Figure 6.**
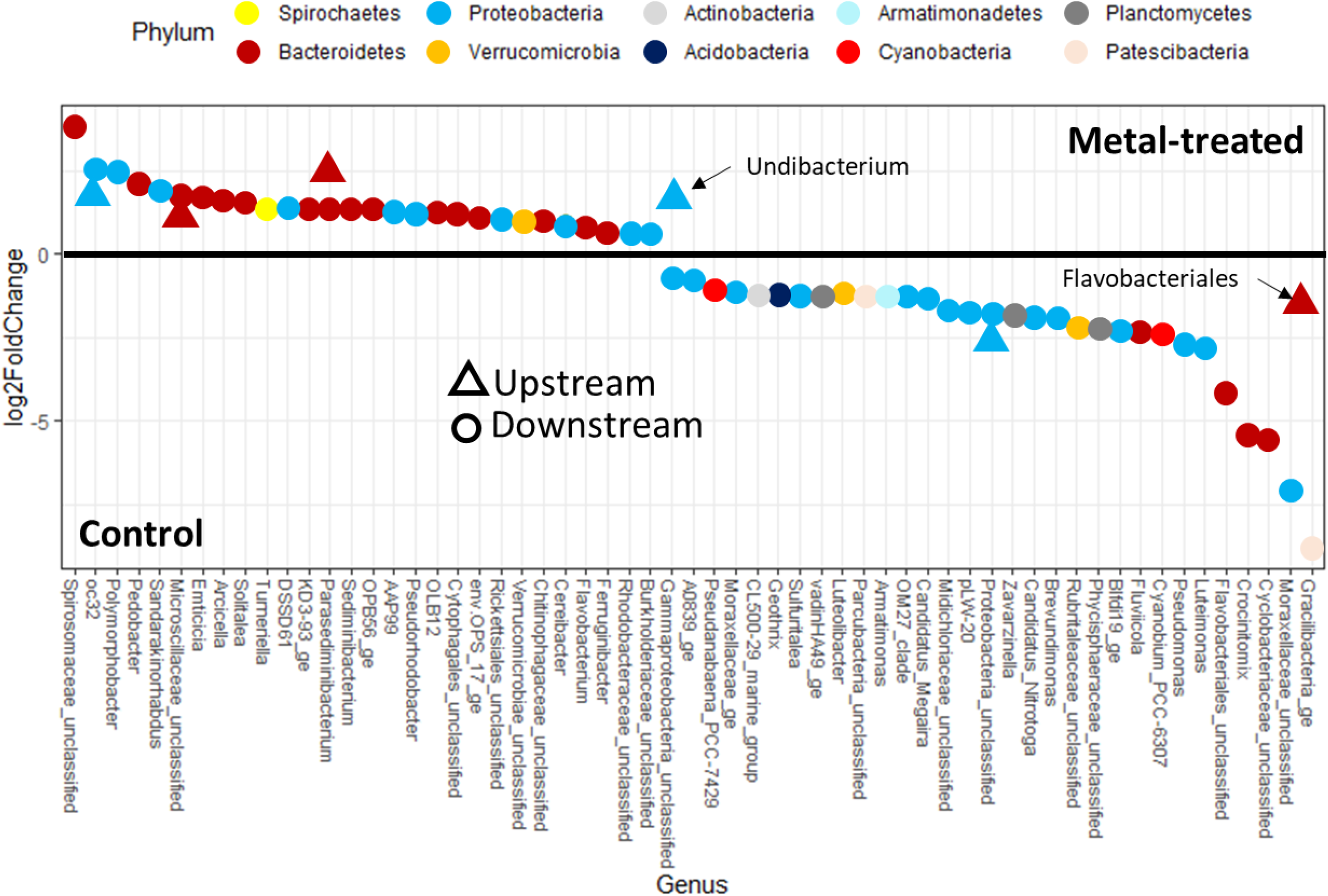
Log2-fold change plot between metal-treated vs. control samples from upstream (triangles) and downstream (circles) communities. Note, Unibacterium *and* Flavobacteriales were drawn in the figure and do not correspond to the taxa listed on the x-axis.

When we compared our observational and mesocosm studies, there were no representative genera that were enriched in downstream sites and metal-treated mesocosms. Additionally, did not find any genera that were enriched from the upstream site in the Upper Arkansas River and in the control mesocosms.

## Discussion

In the observational study of the Upper Arkansas River, measures of microbiome alpha diversity (richness and Shannon Index values) were not different between sites upstream and downstream of California Gulch despite consistently higher sediment metal concentrations of Cd, Cu, and Zn at downstream sites. However, the California Gulch microbiome did have significantly lower richness and Shannon Index values than either the upstream or downstream locations. These results suggest that metal concentrations may need to exceed a certain threshold before having significant effects on microbiome diversity. Previous research has also shown that mining activities or metal impaired streams have very little impact on microbiome diversity, particularly when the metal contamination does not have a pronounced impact on pH (33, 34), as was the case in this study. Similar effects of metal exposure on macroinvertebrate diversity have been reported from these same locations in the Upper Arkansas River. Specifically, overall species richness for macroinvertebrates upstream and downstream of California Gulch were found to be similar, both within and between seasons (17). However, membership for the macroinvertebrate community was different between upstream and downstream locations (18) with the metals disproportionately affecting some taxa more than others.

When we examined membership of the river microbiome at each location, we found significant differences in membership between upstream and downstream locations (Figure 3). Because metal delivery to the stream is most pronounced during Spring run-off, we hypothesized that differences in microbiome membership between locations would be greatest in the springtime, and downstream membership would change the most between seasons. However, seasonal variation in microbiome membership occurred at both upstream and downstream locations. When we examined microbiome membership among sites the majority (39.6%) of the variance in membership explained by the first two principal coordinates was driven by the differences between the California Gulch microbiome and the two main stem locations. Although subtle, membership of the downstream microbiome was more similar to the California Gulch microbiome and this similarity was more pronounced in Spring than Fall, consistent with the idea of metals exerting an influence on the membership of downstream microbiomes. The differences in microbiome membership between upstream and downstream sites (Figure 1) are notable because concentration of metals in the surface water downstream of California Gulch are typically below EPA chronic aquatic life criteria (35).

Previous investigations have shown that experimental metal exposure resulted in much greater effect on composition from macroinvertebrate communities sourced from sites upstream of California Gulch (18). In this mesocosm experiment we expected to see greater changes in microbiome membership of upstream microbiomes in response to the metal treatments, since this site has historically had lower metal exposure and we anticipated the microbes would be more sensitive to metal stress. However, we observed a greater change in downstream microbiome membership in response to metal exposure, in contrast to our expectations. One potential explanation for the differences in treatment effect between locations is that the samples from the downstream location had metal-resistant bacteria present within the microbiome whereas the upstream samples did not or had fewer. Thus, the downstream microbiome had a more rapid response to metal exposure than the upstream community after the 10-day treatment. This mechanism is supported by the lower evenness in samples sourced from the downstream location following metal exposure compared to upstream microbiomes (Table 2), suggesting an enrichment of metal tolerant organisms altered the rank abundance of the downstream microbiome. A recent study examining the effects of the antimicrobial drug Ciprofloxacin also reported more pronounced differences in microbiome membership from experimental exposure to Ciprofloxacin along a gradient of urbanized streams in New York (36). The greatest difference in microbiome membership were observed in stream reaches with the highest ambient concentrations of Ciprofloxacin. We posit that the discrepancy between community responses in macroinvertebrates versus microbiomes in response to metal exposure was due to the relative timescale of our study. For instance, over a 10-day period of metal exposure, observed differences of macroinvertebrate membership are by driven by mortality of the original community, whereas, microbiomes may experience multiple generations during that same time period. Thus, microbiome membership was likely not only altered by mortality but also by enrichment of metal tolerant taxa, which was more pronounced at the downstream compared to the upstream sites.

We also observed differences in the amount of metals retained in microbial biomass between microbiomes sourced from different locations. In the metal-treated samples, the downstream microbiomes had approximately 5-8x greater Cu and Zn compared to the mesocosms with sediments sourced from the upstream site (Figure 7). In contrast, metal concentrations in microbial biomass were very similar between locations in the control treatments. One potential mechanism for this may be increased tolerance of downstream microbiomes through greater production of Extracellular Polymeric Substance (EPS). Stream biofilm EPS can retain metals in proteins, polysaccharides, and humic components (37), and bacterial EPS production can be enhanced in the presence of metals (38). Therefore, microbiome metal-tolerance may further exacerbate metal exposure to higher trophic levels by retaining metals in their biomass. Recent studies have shown that that macroinvertebrates derive much of their metals from diet and not just from aqueous exposure (39-41). Interestingly, in a recent experimental study Zn concentrations in periphyton samples at similar levels as the downstream meta-treated biofilms caused dramatic reduction (> 75%) in mayfly abundance (42). Additionally, metal-resistant populations of oligochaetes in Foundry Cove, New York increased metal exposure to higher trophic levels by production of metal-binding proteins in their tissues (43). If the downstream microbiome did produce more EPS in response to metal exposure (thus retaining more metals) this would suggest a mechanism for how the microbial response to environmental stress (increased EPS production) altered the diets of the next trophic level. Dietary exposure to metals, or decrease in resource quality, provides a mechanism that could explain the differences in macroinvertebrate membership between upstream and downstream locations previously reported for this same location (18).

**Figure 7.**
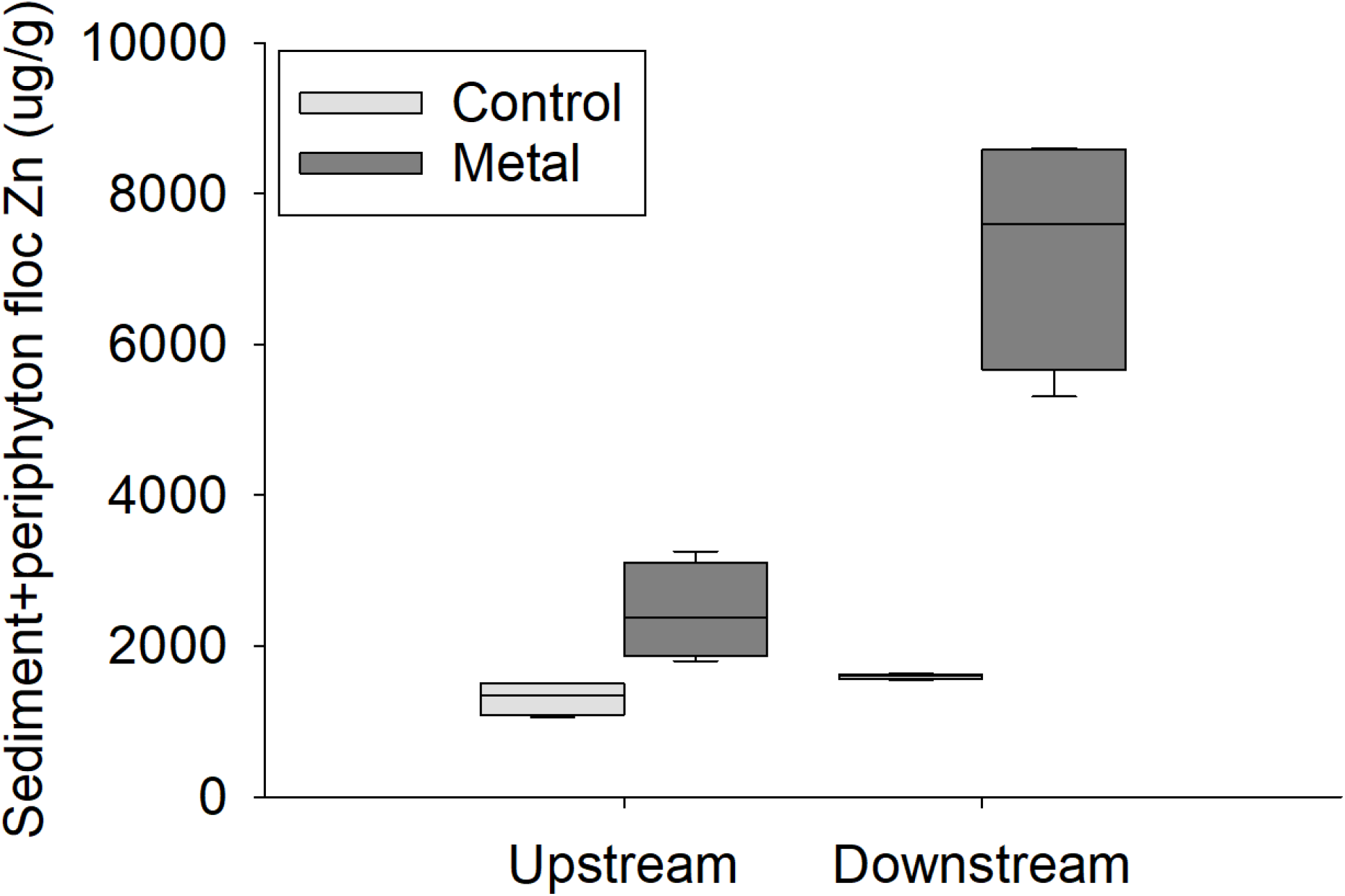
Zinc concentrations of sediment floc after the 10-day mesocosm exposure. All metal-treated samples (“Metals”) were dosed at a target concentration of 650 µg/L Zn.

### Similarity in response between the observational and mesocosm studies

When we compared the membership of microbiomes in the field to those incubated in the experimental streams for ∼10 days we did not observe strong association between samples from observational and mesocosm studies at the genera-level. Whereas the mesocosm experiment was designed to isolate the effect of metals on the stream microbiome our experimental design likely introduced other confounding factors. For example, comparison between the field observations and the mesocosm experiments were complicated by differences in source water chemistry. Mesocosms received natural water inoculum from the hypolimnion of a large reservoir (Horsetooth Reservoir) and not Upper Arkansas River water. Therefore, it is possible that differences in membership between the mesocosm and the field samples may be due in part to differences in source water and the microorganisms that were associated with the water from each ecosystem. In addition, the sites in the Upper Arkansas River are open canopy and the downstream site is located downstream from the town of Leadville, CO (pop. ∼3,000), whereas the reservoir water is comparatively lower in nutrients and sourced from the aphotic hypolimnion. Thus, differences in light environment and water chemistry between the field and the laboratory may also contribute to enrichment of certain genera due to other factors that were not influenced in the same way in the mesocosm studies. Additionally, California Gulch is likely enriched in ammonia and other nutrients from the wastewater treatment process that may be responsible for some observed differences. For instance, we found that genera from the phylum Cyanobacteria were more enriched in the downstream site (i.e., possibly indicating metal-tolerance), but in the mesocosms Cyanobacteria were only significantly enriched in controls. Because we did not see similar enrichment of members of the Cyanobacteria in response to metal exposure in the mesocosm experiments, this would also suggest that differences in Cyanobacteria between locations in the field were not entirely driven by metals but perhaps were due to some of the confounding factors that were present in the observational study.

We observed consistent enrichment of members of a single phylum between both observational and experimental studies and enrichment of members of some genera between both as well. For example, Proteobacteria were prevalent in both upstream and downstream observational samples. This is not surprising given the extremely high diversity within the proteobacteria, but also further highlights the idea that comparisons among different phyla are likely too broad for applied microbial ecology research, as has been previously suggested (44). It is unlikely that metal sensitivity or resistance to metals is a trait that is conserved at the level of the phyla. However, we did find that members of Bacteroidetes were consistently enriched in downstream and metal-treated mesocosms compared to other phyla. This is consistent with previous research that has shown Bacteroidetes to be significantly enriched at other metal polluted sites (45). Three other genera that were significantly enriched in metal-treated mesocosms from both the upstream and downstream locations: *oc32* (Phylum Proteobacteria; Class Gammaproteobacteria; Order Burkholderiales; Family Nitrosomonadaceae), *Parasediminibacterium* (Phylum Bacteroidetes; Class Shingobacteria; Order Sphingobacteriales; Family Chitinophagaceae) and an unclassified genus in the Family Microscillaceae (Phylum Bacteroidetes; Class Cytophagia; Order Cytophagales). Aggregated OTUs from an unclassified genus in the Family Flavobacteriales was found to be enriched in control mesocosms from both upstream and downstream locations. These four genera (e.g., 3 metal-tolerant and 1 metal-sensitive) represent the best candidates from our study to assess the impact of metals on stream ecosystems. Using only the observational study we may have assumed that other genera were also good candidates of metal contamination, however the comparison with the mesocosm experiment excluded these candidates. Since many of the metal-treated genera that were enriched came from the Phylum Bacteroidetes this group may be a logical starting point for more directed research into using microbiome membership as an indicator of metal contamination.

### The effects of low metal exposure to stream ecosystems

The effect of metals on microbiomes upstream and downstream of the contaminant site had similar diversity but significant differences in membership that appeared to be caused by exposure to heavy metals. Other studies have documented changes in microbiome composition associated with a range of heavy metal exposure in the field from diverse geologic sources such as mountaintop mining (33), acid mine drainage (16), and streams influenced by urban runoff (46). In our study, the use of complimentary field observations and experimental mesocosms allowed us to assess which constituents of natural microbiomes were likely to be consistently affected by metal exposure.

We conclude that the microbiomes in the Upper Arkansas River downstream of California Gulch are still responding to metals stress. Even though diversity metrics suggest similarities between upstream and downstream microbiomes, differences in membership indicate that metals are impacting the Upper Arkansas River even after extensive restoration efforts have lowered surface water metals below current US EPA criteria levels (35). Interestingly, typical metal concentrations downstream of California Gulch are not directly toxic to macroinvertebrate communities (47). However, the patterns we observed between upstream and downstream microbiomes (i.e., comparable alpha diversity but distinct membership) were similar to patterns found for the macroinvertebrate communities at the same site (18). We also note that in the mesocosm study the microbiomes sourced from the downstream site accumulated more metals during the experiments than those sourced from the upstream site. We did not see a similar difference in metal content between the upstream and downstream sourced mesocosms for the control experiment. Taken together these results suggest that dietary exposure to metals or changes in microbial biomass that decrease nutritive quality (e.g. generation of excess EPS to metal exposure (48-50), or both may cause shifts in the macroinvertebrate community composition). This mechanism is further supported by the much lower abundance of functional feeding groups in the downstream communities that would be indicative of a dietary shift from biofilm to seston feeders. A previous study (18) found that upstream macroinvertebrates were enriched in mayflies and other “scrapers” (scraper is the common name given to insects of the functional feeding type that “scrape” biofilms from rocks as a food source) and downstream communities were enriched in Caddisflies and other seston-feeding taxa.

### Conclusions

In this current study we cannot conclusively link the response of the microbiome to metals to changes in diet quality of their primary consumers, stream macroinvertebrates. However, several results presented in this study are consistent with the idea that a microbial response to metals at the base of the food web may be affecting consumers one trophic level above. If this is indeed the case, then it suggests that the current criteria that uses chronic exposure of aquatic macroinvertebrates to assess stream health (a threshold below which is thought to be protective of ∼95% of the aquatic community) is insufficient to assess the impact of metals on stream ecosystems. It is becoming increasingly evident that dietary exposure is as important as direct exposure to aquatic life (51-53) and should be considered when assessing the impact of metals on stream ecosystems. One challenge presented here is that quantification of the metal content of macroinvertebrate diets is much harder to measure than the metal content of the surface water. However, if the dietary exposure is mediated through shifts in the microbiome, then microbial metrics, as presented here, may provide a better alternative to assess the impact of metals on stream ecosystems. Our research suggests current best practice guidelines of stream water quality (e.g., EPA aquatic life criteria) may miss impacts of metal contamination on the community that form the base of the stream ecosystems and additional factors (e.g., dietary exposure, microbial metrics) should be included as these standards are improved.

## Acknowledgements

Portions of this work were supported by NSF IOS award DEB#1456959 awarded to EKH, the Warner College of Natural Resources, the Colorado Water Center, and the Colorado Mountain Club, Colorado Water Institute, and SETAC Student Training Exchange Opportunity awards to BAW. We thank Colorado State University Natural Resource Ecology Laboratory and Aquatic Ecotoxicology Laboratory students and staff for laboratory assistance.

